# STAT5B leukemic mutations, altering SH2 tyrosine 665, have opposing impacts on immune gene programs

**DOI:** 10.1101/2024.12.20.629685

**Authors:** Hye Kyung Lee, Jichun Chen, Rachael L. Philips, Sung-Gwon Lee, Xingmin Feng, Zhijie Wu, Chengyu Liu, Aaron B. Schultz, Molly Dalzell, Foster Birnbaum, Joel A. Sexton, Amy E. Keating, John J. O’Shea, Neal S. Young, Alejandro V. Villarino, Priscilla A. Furth, Lothar Hennighausen

**Affiliations:** Laboratory of Genetics and Physiology, National Institute of Diabetes and Digestive and Kidney Diseases, US National Institutes of Health, Bethesda, Maryland 20892, USA; Hematology Branch, National Heart, Lung, and Blood Institute, National Institutes of Health, Bethesda, Maryland 20892, USA; Molecular Immunology and Inflammation Branch, National Institute of Arthritis and Musculoskeletal and Skin Diseases, National Institutes of Health, Bethesda, Maryland 20892, USA; Transgenic Core, National Heart, Lung, and Blood Institute, US National Institutes of Health, Bethesda, Maryland 20892, USA; Department of Microbiology and Immunology, Miller School of Medicine, University of Miami, Miami, Florida, 33146 USA; Sylvester Comprehensive Cancer Center, University of Miami, Miami, Florida, 33146 USA; Department of Biology, Massachusetts Institute of Technology, Cambridge, Massachusetts, 02139, USA; Comutational and Systems Biology, Massachusetts Institute of Technology, Cambridge, Massachusetts, 02139, USA; Department of Biological Engineering, Massachusetts Institute of Technology, Cambridge, Massachusetts, 02139, USA; Koch Institute for Integrative Cancer Research, Massachusetts Institute of Technology, Cambridge, Massachusetts, 02139, USA

## Abstract

STAT5B is a vital transcription factor for lymphocytes. Here, function of two STAT5B mutations from human T cell leukemias: one substituting tyrosine 665 with phenylalanine (STAT5B^Y665F^), the other with histidine (STAT5B^Y665H^) was interrogated. *In silico* modeling predicted divergent energetic effects on homodimerization with a range of pathogenicity. In primary T cells *in vitro* STAT5B^Y665F^ showed gain-of-function while STAT5B^Y665H^ demonstrated loss-of-function. Introducing the mutation into the mouse genome illustrated that the gain-of-function *Stat5b*^Y665F^ mutation resulted in accumulation of CD8^+^ effector and memory and CD4^+^ regulatory T-cells, altering CD8^+^/CD4^+^ ratios. In contrast, STAT5B^Y665H^ ‘knock-in’ mice showed diminished CD8^+^ effector and memory and CD4^+^ regulatory T cells. In contrast to wild-type STAT5, the STAT5B^Y665F^ variant displayed greater STAT5 phosphorylation, DNA binding and transcriptional activity following cytokine activation while the STAT5B^Y665H^ variant resembled a null. The work exemplifies how joining *in silico* and *in vivo* studies of single nucleotides deepens our understanding of disease-associated variants, revealing structural determinants of altered function, defining mechanistic roles, and, specifically here, identifying a gain-of function variant that does not directly induce hematopoietic malignancy.

## Introduction

Transcription factors belonging to the Signal Transducers and Activators of Transcription (STAT) family are activated by upstream cytokines and growth factors and, in turn, regulate both universal and lineage-specific genetic programs (Philips *et al*, 2022). STAT5A (Liu *et al*, 1995; Wakao *et al*, 1994) and STAT5B (Azam *et al*, 1995; Liu *et al*., 1995; Mui *et al*, 1995), ancestral members of the STAT family, play defining roles within the hematopoietic system (Yao *et al*, 2006), the mammary gland (Cui *et al*, 2004; Liu *et al*, 1997; Miyoshi *et al*, 2001), body growth (Davey *et al*, 1999; Holloway *et al*, 2007; Hwa, 2021), and liver metabolism (Holloway *et al*., 2007). As ‘signal-dependent’ Transcription Factors (TFs), they instruct common and lineage-restricted enhancers and super-enhancers (Lee *et al*, 2023; Li *et al*, 2017; Shin *et al*, 2016; Villarino *et al*, 2016; Villarino *et al*, 2017; Yamaji *et al*, 2013), thereby controlling a wide range of genetic programs, in tissues as diverse as the mammary gland, liver and the immune system.

While genetic programs regulated by native STAT5A and STAT5B are well investigated, the impact of missense mutations on these TFs has not been studied in detail. Based on current literature and genomic databases, at least one third of the amino acids of STAT5 can acquire germline and somatic missense mutations in humans. Inactivating STAT5B germline mutations (Bhattacharya *et al*, 2022; Hwa, 2021; Rajala *et al*, 2013) have been associated with GH insensitivity (Laron syndrome) and immune pathology, while activating somatic mutations (Bhattacharya *et al*., 2022; Kiel *et al*, 2014; Kontro *et al*, 2014; Rajala *et al*., 2013) have been identified in patients with T cell large granular lymphoblastic leukemia (T-LGLL) (Bhattacharya *et al*., 2022; Kiel *et al*., 2014; Kontro *et al*., 2014; Rajala *et al*., 2013) and T-cell prolymphocytic leukemia (T-PLL) (Kiel *et al*., 2014). T-cell large granular lymphocytic leukemia (T-LGLL) is a rare disorder marked by the clonal expansion of cytotoxic T cells within the peripheral blood and bone marrow (Matutes, 2018). Among the two subtypes of T-LGLL, the more frequent CD8 T-LGLL is characterized by STAT3 mutations while the CD4 T-LGLL has been linked to mutations in the STAT5B, that are clustered in the SH2 and C-terminal transactivation domain. Accordingly, N642H and Y665F are the most frequent STAT5B mutations, and both lead to increased STAT5 activity (Andersson *et al*, 2016; Bhattacharya *et al*., 2022). The SH2 domain is central to cytokine-induced, JAK-dependent STAT5B tyrosine phosphorylation, activation, dimerization, nuclear translocation and establishment of functional transcription enhancers controlling genetic programs. A total of six somatic missense mutations and four germline mutations within the SH2 domain of STAT5B have been reported in the literature (Bhattacharya *et al*., 2022; Hwa, 2021). While cell culture experiments have established their capacity to function as GOF or LOF, their complex physiological and genomic impacts have not been investigated. Of particular interest are two Single Nucleotide Variants (SNV) identified in patients with T cell Large Granular Lymphocytic Leukemia (T-LGLL) and T-PLL (Kontro *et al*., 2014). Both are found within the SH2 domain and change tyrosine 665 (Y665) to either phenylalanine (Y665F) or histidine (Y665H) (Kiel *et al*., 2014; Rajala *et al*., 2013). Both mutations have been described as Gain-of-Function (GOF) *in vitro* (Pham *et al*, 2018), suggesting that they activate similar and possibly overlapping genetic programs. To determine if they are driver mutations for T cell leukemias, we integrated STAT5B^Y665F^ and STAT5B^Y665H^ into the mouse genome and examined their impact on lymphocyte development and function. Using this approach, we show that only the STAT5B^Y665F^ mutation displays GOF characteristics *in vivo* while STAT5B^Y665H^ is a Loss-Of-Function (LOF) variant. Significantly neither directly induced malignant transformation. These data expand our understanding of the structural and molecular impact of these two specific single nucleotide STAT5B variants in lymphocyte homeostasis and function, identifying specific genes and pathways involved in their pathogenesis.

## Results

### Structural impact of the STAT5B^Y665F^ and STAT5B^Y665H^ missense mutations *in silico*

Tyrosine 665 is the second most abundant mutational target in the STAT5B SH2 domain (Fig. 1A), with 12 T cell leukemia cases reported in the COSMIC database and 15 cases in the database of the Munich Leukemia Laboratory (MLL). To gauge the impact of the STAT5B^Y665F^ and STAT5B^Y665H^ mutations, we conducted comprehensive *in silico* analyses, combining structural prediction and pathogenicity assessment tools. Using AlphaFold3, we generated structures of the STAT5A and STAT5B SH2 domain homodimers. As expected from their highly conserved sequences, STAT5A and STAT5B produced nearly identical structure predictions by AlphaFold3 (Abramson *et al*, 2024) (Supplementary Fig. 1). The AlphaFold3-predicted structure of the STAT5B SH2 domain homodimer revealed that STAT5B^Y665^ is located at a critical interface involved in STAT5B homo-dimerization (Fig. 1B). This residue position and its surrounding structural elements are highly similar to what is predicted in the model of the STAT5A dimer interface generated by Fahrenkamp et al. (Fahrenkamp *et al*, 2016).

**Fig. 1.**
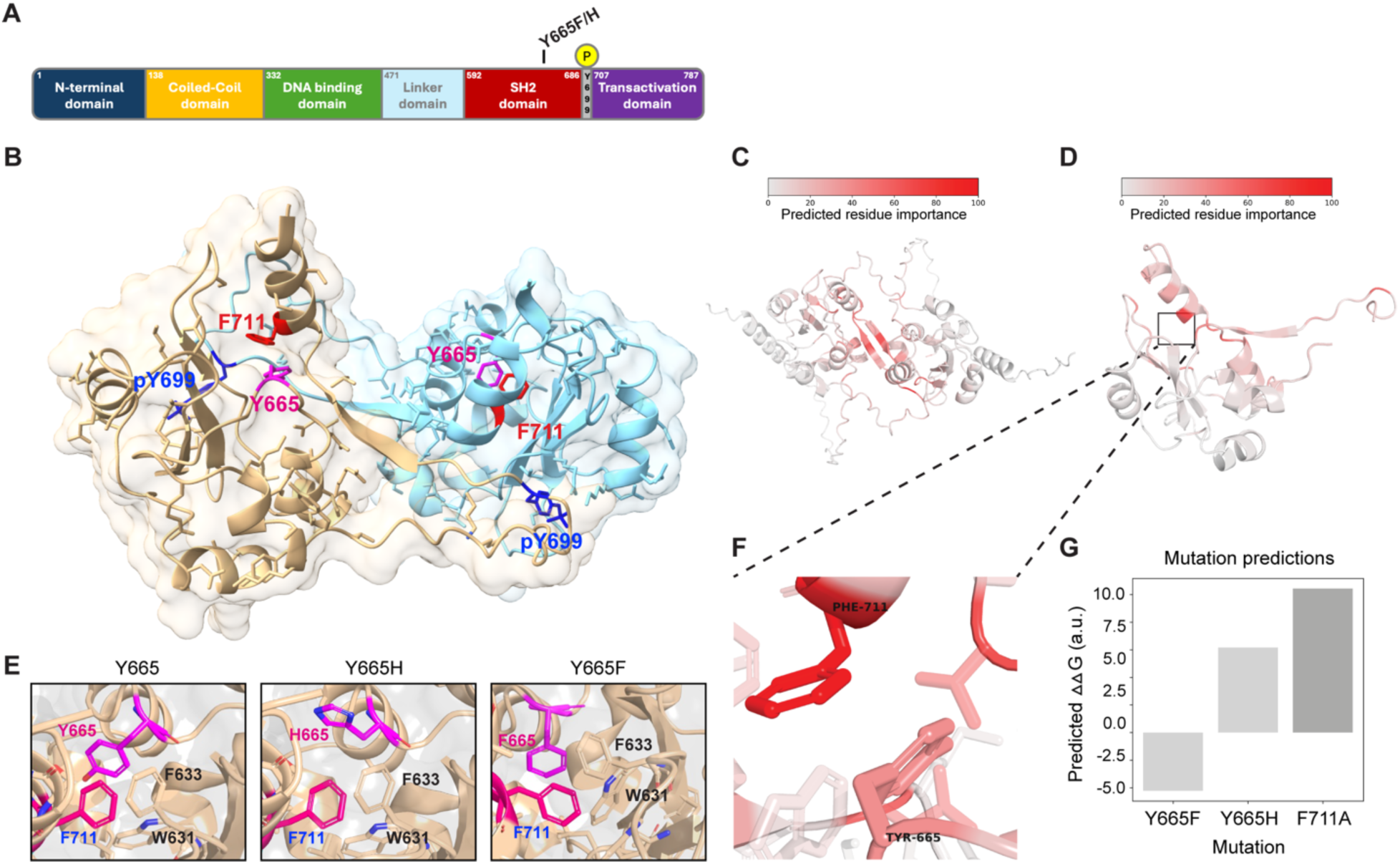
STAT5B SH2 dimerization modeled by AlphaFold3. **A** Schematic of human STAT5B protein domains showing the locations of tyrosine 665 (Y665) and phospho-tyrosine 699 (pY699). **B** Structure of the human STAT5B SH2 homodimer generated by AlphaFold3. Binding pockets of key residues pY699 (blue) and F711 (red) are indicated. The STAT5B model shows the hydrophobic binding pocket containing key residue Y665 (magenta). **C** Structure of the human STAT5B SH2 homodimer generated by AlphaFold3 with residues colored red relative to their importance to the binding interface as predicted by COORDinator. **D** Structure of the human STAT5B SH2 monomer in its dimeric conformation generated by AlphaFold3 with residues colored red relative to their importance to stabilization of the C-terminal tail as predicted by COORDinator. **E** Model of the human STAT5B SH2 homodimer with tyrosine, histidine, or phenylalanine at position 665, as predicted by AlphaFold3. **F** The STAT5B model highlights the intramolecular interaction between F711 and the hydrophobic binding pocket containing key residue Y665. **G** Comparison of the predicted energetic consequences of the F711A, Y665F, and Y665H mutations on stabilization of the C-terminal tail. Relative mutational effects are annotated using ΔΔG, in arbitrary units; stabilizing mutations have a negative value, and destabilizing mutations have a positive value.

We predicted the energetic contribution of each residue in the structures of both STAT5A and STAT5B homodimers using COORDinator that is a neural network trained to take as input a protein backbone structure and output a sequence compatible with that backbone (Li *et al*, 2023). The version of COORDinator that we used was fine-tuned to predict the effects of amino-acid substitutions on protein stability (Tsuboyama *et al*, 2023) using only the backbone structure as input. To distinguish energetic contributions specific to dimerization from more general contributions to domain stability, we compared the predicted energies of the STAT5B and STAT5A SH2 domain residues with and without their homodimeric counterparts (Supplementary Fig. 2A-B). This approach highlighted key residues within the C-terminal tail that form the antiparallel dimeric interface, as well as amino acids C-terminal to the phosphorylated Y699 in STAT5B (Fig. 1C-D). We also used COORDinator to predict the energies of intramolecular interactions between the STAT5B C-terminal tail and the SH2 domain (Supplementary Fig. 2C). This approach highlighted STAT5B^F711^ as making the greatest contribution to stabilizing the intramolecular interactions that support the dimer conformation (Fig. 1E-F). Most substitutions to STAT5B^F711^ and interacting residues such as STAT5B^Y665^ were predicted to destabilize the intramolecular interaction (Fig. 1G). Importantly, the STAT5B^Y665H^ substitution introduces an imidazole and was predicted by COORDinator to destabilize binding of the C-terminal tail (Fig. 1G). In contrast, STAT5B^Y665F^ was predicted to stabilize the structure, perhaps by promoting intramolecular aromatic stacking interactions with STAT5B^F711^.

To further evaluate the potential pathogenicity of these two mutations, we employed multiple state-of-the-art computational prediction tools. AlphaMissense (Cheng *et al*, 2023b) predicted mild functional impact for both mutations with scores of 0.173 and 0.383 for STAT5B^Y665F^ and STAT5B^Y665H^ respectively, categorizing them as benign. However, the Combined Annotation Dependent Depletion (CADD) (Rentzsch *et al*, 2019) PHRED scores of 24.3 (STAT5B^Y665F^) and 23.1 (STAT5B^Y665H^) suggested potential deleterious effects, as scores above 20 are often considered impactful. REVEL (Rare Exome Variant Ensemble Learner) (Ioannidis *et al*, 2016) analysis yielded scores of 0.535 for STAT5B^Y665F^ and 0.304 for STAT5B^Y665H^, indicating a higher probability of pathogenicity for STAT5B^Y665F^. Notably, PolyPhen-2 (Adzhubei *et al*, 2013b) showed marked differences between the mutations, with STAT5B^Y665F^ scoring 0.93 (probably damaging) and STAT5B^Y665H^ scoring 0.084 (benign). In summary, different tools provided a range of pathophysiological predictions for the mutants. Only COORDinator, the only tool that uses structural modeling to predict energetic effects rather than pathogenicity, gave predictions that fully match the experimental results.

### STAT5B^Y665F^ but not STAT5B^Y665H^ activates genetic programs in CD4^+^ T cells *in vitro*

To assess the impact of the STAT5B^Y665F^ and STAT5B^Y665H^ mutations on the transcriptional activity of STAT5B, we measured gene expression in *Stat5a/b*^-/-^ T cells reconstituted with either wild-type or mutant STAT5B. Naïve T cells were purified from lymph nodes and spleens of wild-type or *Stat5a/b*^-/-^ mice (Cui *et al*., 2004), activated *in vitro* and transduced with retroviral vectors encoding wild-type *STAT5B*, STAT5B^Y665F^ and STAT5B^Y665H^ (Fig. 2A). STAT5B^N642H^, an established activating mutation identified in T cell leukemia patients (de Araujo *et al*, 2019), was included as a positive control. We confirmed an approximately 98% reduction of *Stat5b* mRNA in *Stat5a/b*^-/-^ CD4^+^ cells as compared to wild-type cells, demonstrating efficient deletion of the *Stat5a/b* locus and providing a null background for testing the different mutants (Fig. 2B). Upon retroviral introduction of the various constructs, *Stat5b* mRNA levels were restored to near WT levels in the *Stat5a/b*^-/-^ cells. Notably, both positive control STAT5B^N642H^ and experimental STAT5B^Y665F^ induced surface expression of the two well-known STAT5 target genes CD25 and CD98 (Villarino *et al*, 2022), while STAT5B^Y665H^ failed to do so (Fig. 2C). Transcriptome analysis showed that wild-type STAT5B mobilized 141 Differentially Expressed Genes (DEGs) versus ‘empty’ retrovirus, while positive control STAT5B^N642H^ and experimental STAT5B^Y665F^ and STAT5B^Y665H^ induced 363, 318 and 23 genes, respectively (Supplementary Table 1). Of the 318 genes mobilized by STAT5B^Y665F^, about one third (113 genes) are directly engaged by STAT5B as determined by ChIP-seq and are therefore considered *bona fide* STAT5B target genes (Supplementary Table 1). Many DEGs (287) were shared between the positive control STAT5B^N642H^ and experimental STAT5B^Y665F^ mutants, indicating that, overall, these variants demonstrated similar transcriptional effects (Fig. 2D; Supplementary Table 1). Common *bona fide* STAT5 target genes, such as *Socs2* and *Cish*, were activated up to several thousand-fold, while more cell-restricted genes, such as lymphotoxin alpha (*Lta*), *Irf8* and the interleukin 2 receptor alpha (*Il2ra*) were induced up to 10-fold (Fig. 2E-G). In contrast, STAT5B^Y665H^ failed to activate gene expression in the *Stat5a/b*^-/-^ background. These results demonstrate that STAT5B^Y665F^ and STAT5B^N642H^ have equivalent Gain-Of-Function (GOF) capacity, while STAT5B^Y665H^ is a clear Loss-Of-Function (LOF) mutation.

**Fig. 2.**
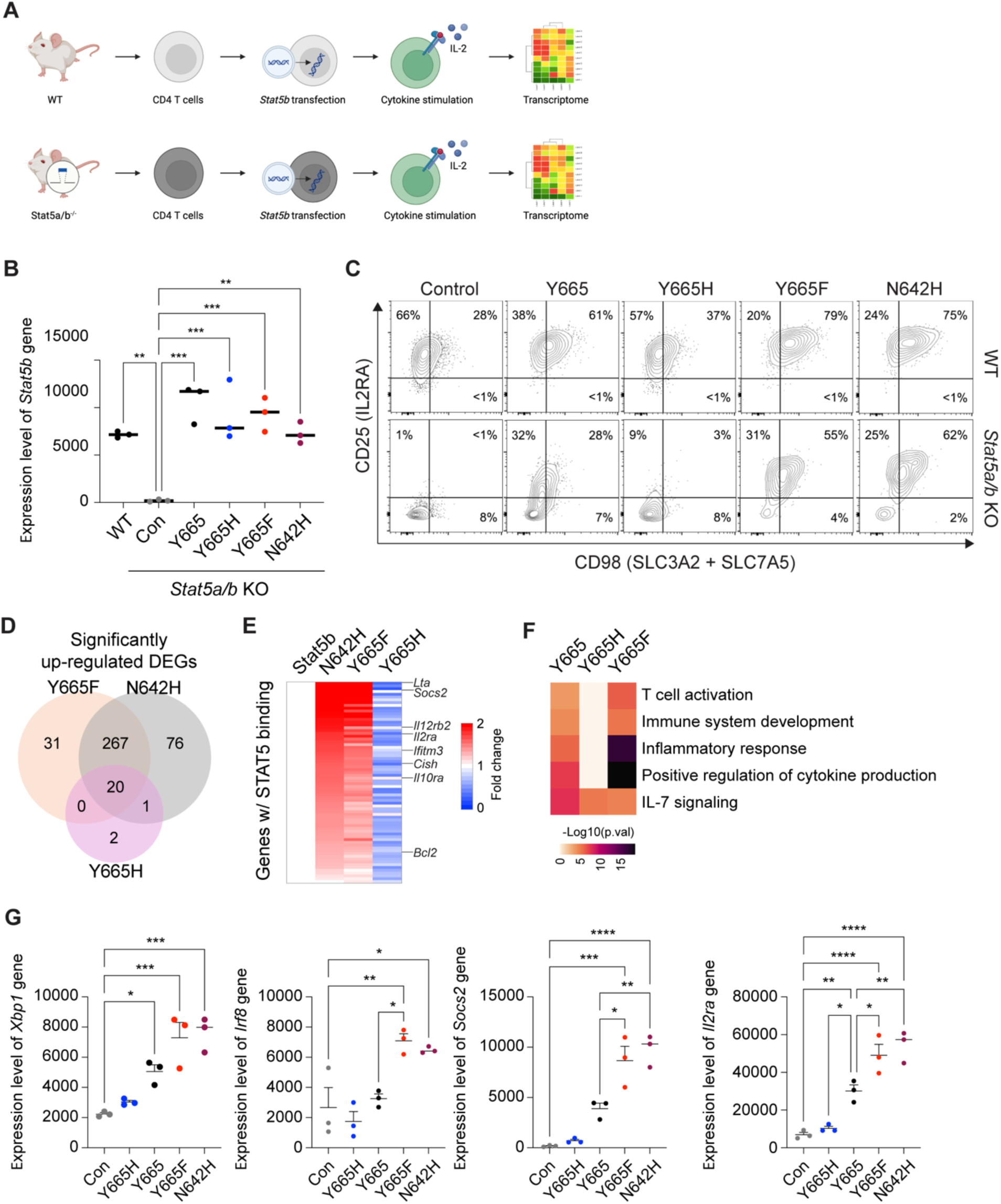
Transcriptomes activated by wild-type and STAT5B mutations in *Stat5a/b*-null T cells. **A** Schematic of the experimental approach to assess the impact of STAT5B^Y665^ mutations on naïve CD4 T cells from lymph nodes and spleens of wild-type or STAT5A/B-deficient mice. **B** The expression level of *Stat5b* gene transduced with retrovirus encoding STAT5B wild-type Y665, Y665F or Y665H. **C** Flow cytometry contour plots showing CD4 cell population (*n* = 3). **D** Venn diagram displaying the number of significantly induced genes by retroviral vectors encoding Y665F, Y665H and N642H mutations compared to wild type. Cells were stimulated with IL-2 (*n* = 3). **E** Heatmaps showing fold changes of significantly upregulated genes between N642H, Y665F and Y665H on Stat5b (Y665) as 1 in STAT5A/B deficient T cells. **F** Heatmap of genes expressed at significantly higher levels in each sample and significantly enriched in Gene Ontology (GO) terms. **G** Dot plots of the normalized read counts for mRNA levels of five genes regulated by STAT5B. Results are shown as the means ± SEM of independent biological replicates. *P*-value are from one-way ANOVA with Tukey’s multiple comparisons test. \**P* < 0.05, \*\**P* < 0.01, \*\*\**P* < 0.0001, \*\*\*\**P* < 0.0001.

### Introduction of *Stat5b*^Y665F^ and *Stat5b*^Y665H^ into the mouse genome

To investigate *in vivo* consequences of the STAT5B^Y665F^ and STAT5B^Y665H^ mutations, we introduced them into mice genome using CRISPR/Cas9 and base editing (see M&M for details). Of note, while the expected 25% homozygous was obtained from *Stat5b*^Y665H^ heterozygous mating, only 13% homozygosity was obtained by breeding heterozygous *Stat5b*^Y665F^ mice (Lee *et al*, 2024). The reason for the loss of approximately 50% of the homozygous mice is not known. Both *ex vivo* and *in vivo* studies were performed (Fig. 3A). While homozygous *Stat5b*^Y665F^ and *Stat5b*^Y665H^ mice displayed an overall normal appearance at the time of weaning, the latter had a retarded body growth, consistent with suboptimal growth hormone signaling in the presence of a LOF Stat5b mutation. An especially progressive and morbid form of ulcerative dermatitis, reminiscent of the type known to occur in the inbred C57*Bl/*6 strain, developed in the *Stat5b*^Y665F^ mice necessitating humane euthanasia of both heterozygous and homozygous mice at early ages. Notably, the condition developed at a significantly earlier age in the homozygous as compared to heterozygous mice. (Fig. 3B). In contrast, homozygous *Stat5b*^Y665H^ mice did not develop either dermatitis or any other condition requiring early euthanasia over the first year of life. Both heterozygous and homozygous *Stat5b*^Y665F^ mice demonstrated splenomegaly as compared to wild-type mice while significantly smaller spleens were observed in homozygous *Stat5b*^Y665H^ mice (Fig. 3C).

**Fig. 3.**
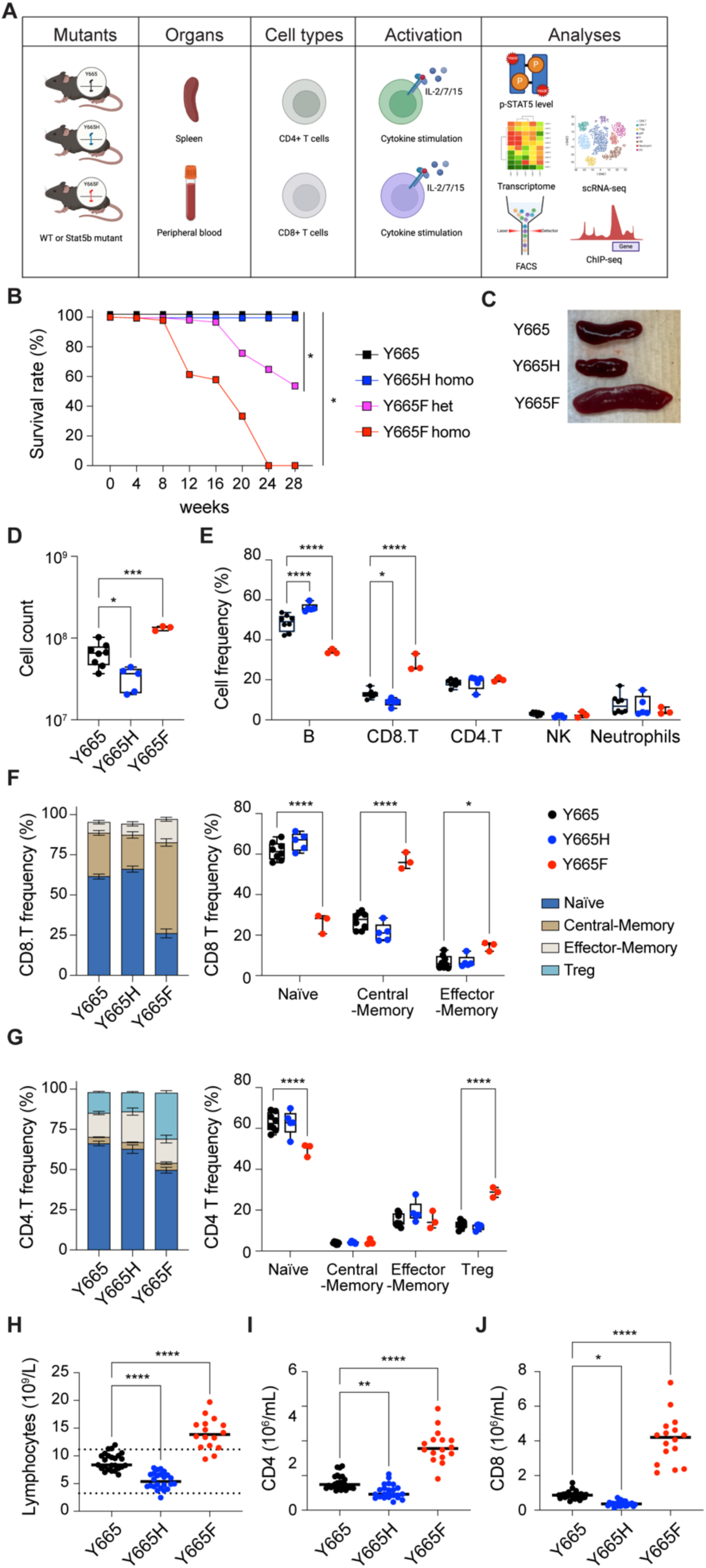
Altered T cell distribution in spleens of *Stat5b* mutant mice. **A** Schematic illustration of the experimental approach. The Stat5^Y665^ variants were introduced into mouse genome, and T cells and stem cells from immune system organs. Intact or cytokine stimulated cells were subjected to western blot, RNA-seq, scRNA-seq, FACS and ChIP-seq analyses. **B** Kaplan-Meier curves exhibiting the survival rate. Y665, *n* = 101; Y665H, *n* = 187; Y665F^+/-^, *n* = 217; Y665F^-/-^, *n* = 46. **C** Images of spleens from WT and mutant mice. **D** Box plots showing total cell number in spleen from wild-type and mutant mice analyzed via flow cytometry (Y665, *n* = 8; Y665H, *n* = 5; Y665F, *n* = 3). *P*-value are from one-way ANOVA with Tukey’s multiple comparisons test. Median, middle bar inside the box; IQR, 50% of the data; whiskers, 1.5 times the IQR. **E** Percentages of B cells, CD4 T cells, CD8 T cells, NK cells and neutrophil were calculated using FACS. **F-G** Percentages of CD8 and CD4 subpopulations. *P*-value are from two-way ANOVA with Tukey’s multiple comparisons test. \**P* < 0.05, \*\**P* < 0.01, \*\*\**P* < 0.0001, \*\*\*\**P* < 0.0001. **H** Count of lymphocytes in peripheral blood from 7-10-week-old adult wild-type and mutant mice. Results are shown as the median of independent biological replicates (Y665, *n* = 25; Y665H, *n* = 24; Y665F, *n* = 16). **I and J** Numbers of CD4+ and CD8+ T cells identified by flow cytometry. Results are shown as the median of independent biological replicates. Statistical significance was assessed using one-way ANOVA followed by Tukey’s multiple comparisons test.

### Increased CD8^+^ central memory cells and CD4^+^ Tregs in splenocytes from *Stat5b*^Y665F^ mice

To address the question of which cell populations were present in the differentially sized spleens of the *Stat5b*^Y665F^ and *Stat5b*^Y665H^ mice, we quantified splenic immune cell populations, focusing on effector, regulatory, and memory T cells using flow cytometry (Fig. 3D-G). Homozygous *Stat5b*^Y665F^ mice exhibited a significantly increased total splenocyte number, whereas *Stat5b^Y665H^*mice displayed a significant reduction in cell numbers as compared to wild-type controls (Fig. 3D). Total CD8^+^ cells were expanded in *Stat5b*^Y665F^ mice but significantly reduced in *Stat5b*^Y665H^ mice (Fig. 3E). No significant differences were observed between the two mutants in CD4^+^ and neutrophil populations, and natural killer (NK) cells. Central and effector memory cells, crucial for long-term immunological memory, immune surveillance, and response within peripheral tissues, were significantly and selectively elevated within the CD8^+^ T cell compartment in the *Stat5b*^Y665F^ mice (Fig. 3F). In contrast these findings were not present in the CD4^+^ T cell compartment. Regulatory T cells (Tregs) showed extensive expansion in *Stat5b*^Y665F^ mice (Fig. 3G), while reduced STAT5B function in *Stat5b*^Y665H^ mice was not associated with significant changes in Treg numbers. We then investigated whether or not similar changes in T cell compartments could be identified in peripheral blood. Similar to the spleen, we noted a significant increase of lymphocytes, CD8^+^ and CD4^+^ T cells in two months-old homozygous *Stat5b*^Y665F^ mice and a significant decrease in *Stat5b*^Y665H^ mice (Fig. 3H-J). To test whether or not these differences were stable with age, repeat assays were performed on 11-month-old mice. At that timepoint, due to the necessity for humane euthanasia at earlier ages in the *Stat5b*^Y665F^ homozygous mice, only heterozygous *Stat5b^Y665F^* mice were available for comparison with the homozygous *Stat5b*^Y665H^ mice. Findings in older heterozygous *Stat5b*^Y665F^ mice resembled those found in younger homozygous *Stat5b*^Y665F^ mice and no significant differences between younger and older *Stat5b*^Y665H^ mice were found. (Supplementary Fig. 3-4). Notably, no leukemic cells were identified in either the *Stat5b*^Y665F^ or *Stat5b*^Y665H^ mice at either young or older age.

### Expansion of central memory T cells in *Stat5b*^Y665F^ mice

To further investigate the effects of mutant Stat5 on gene expression in T cells, we isolated CD45^+^ cells from spleens of wild type and mutant mice and performed single-cell RNA sequencing (scRNA-seq). CD45^+^ is a major transmembrane glycoprotein expressed on all nucleated hematopoietic cell and a well-established white blood cell marker. After identifying key T cell populations (Fig. 4A), we noted that total CD8^+^ cells were expanded in *Stat5b*^Y665F^ mice and reduced in *Stat5b*^Y665H^ mice, as were B cells and monocytes (Fig. 4B). Central memory T cells (Tcm) were significantly expanded in Stat5b^Y665F^ mice compared to wild-type or Stat5b^Y665H^ mice, similar to the flow cytometry results (Fig. 4C). Known Stat5-regulated genes were also enriched in *Stat5b*^Y665F^ Tcm including *Gzmb*, *Cish* and *Il12rb*, (Fig. 4D). The scRNA-seq also revealed an expansion of CD4^+^ T_regs_ exclusively in *Stat5b*^Y665F^ mice (Fig. 4E), with expression of Stat5-dependent genes associated with cellular homeostasis, Th17 cell differentiation and inflammatory responses in these cells (Fig. 4F, G). These findings highlight the distinct roles of activating and inactivating STAT5B mutations in shaping T cell subtypes and their functional states

**Fig. 4.**
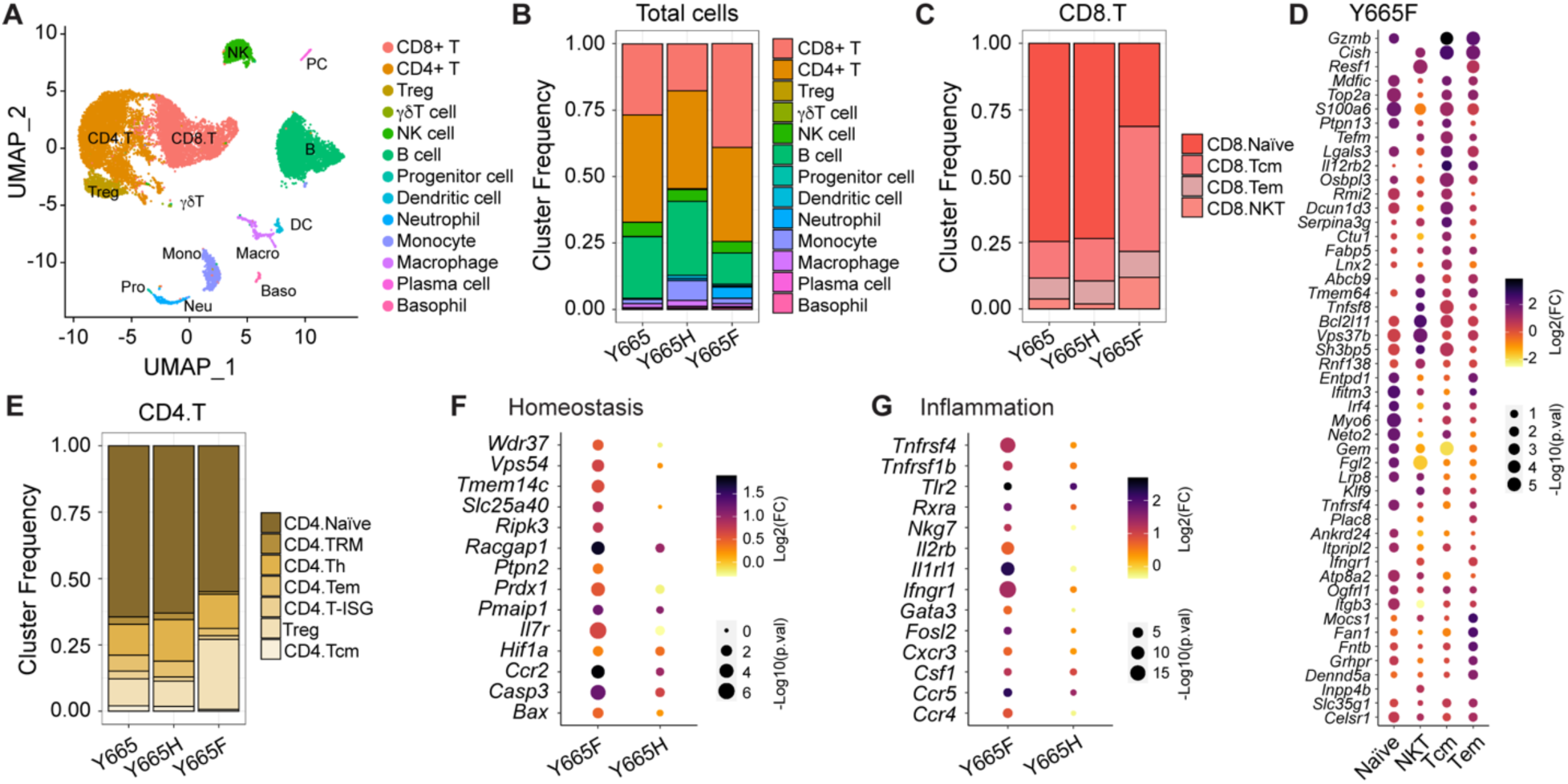
Immune dynamics in spleen analyzed using scRNA-seq. **A** UMAP projection of lymph node from WT and mutant mice, colored by cell population (*n* = 1 combined sample from three mice of each group). **B** Heatmap of cell population. **C** Frequencies of CD8 subtypes: Tcm (central memory T cells), Tem (effector memory T cells), and NKT (NK-like T cells). **D** Dot plot of differentially expressed genes related to cytokine signaling and T cell activation in CD8 T cell subsets. **E** Frequencies of CD4 subtypes: TRM (tissue resident memory T cells), Th (helper T cells), and Treg (regulatory T cells). **F-G** Dot plot of differentially expressed genes related to homeostasis (F) and inflammatory response (G) in regulatory CD4 T cell subpopulations.

### STAT5B^Y665F^ demonstrates enhanced tyrosine phosphorylation of upon interleukin stimulation

Tyrosine phosphorylation of both STAT5A and STAT5B is required for transcriptional capacity and immune cell regulation (Lin *et al*, 2024). To gauge the impact of two STAT5B mutations, we cultured spleen-derived primary T cells from wild-type and mutant mice with IL-2, IL-7 and IL-15, and measured phospho-STAT5B (p-STAT5B) by flow cytometry (Fig. 5A-B). All three interleukins induced tyrosine phosphorylation of STAT5B^Y665F^ but not STAT5B^Y665H^ in CD4^+^ and CD8^+^ T cells. Importantly, phosphorylation levels were significantly higher in stimulated *Stat5b*^Y665F^ cells compared to stimulated wild-type controls. This difference was most prominent in CD4^+^ cells, where p-STAT5B^Y665F^ levels exceeded wild type levels by 5-10-fold (Fig. 5A-B). These results support the notion that STAT5B^Y665F^ and STAT5B^Y665H^ represent GOF and LOF mutants, respectively.

**Fig. 5.**
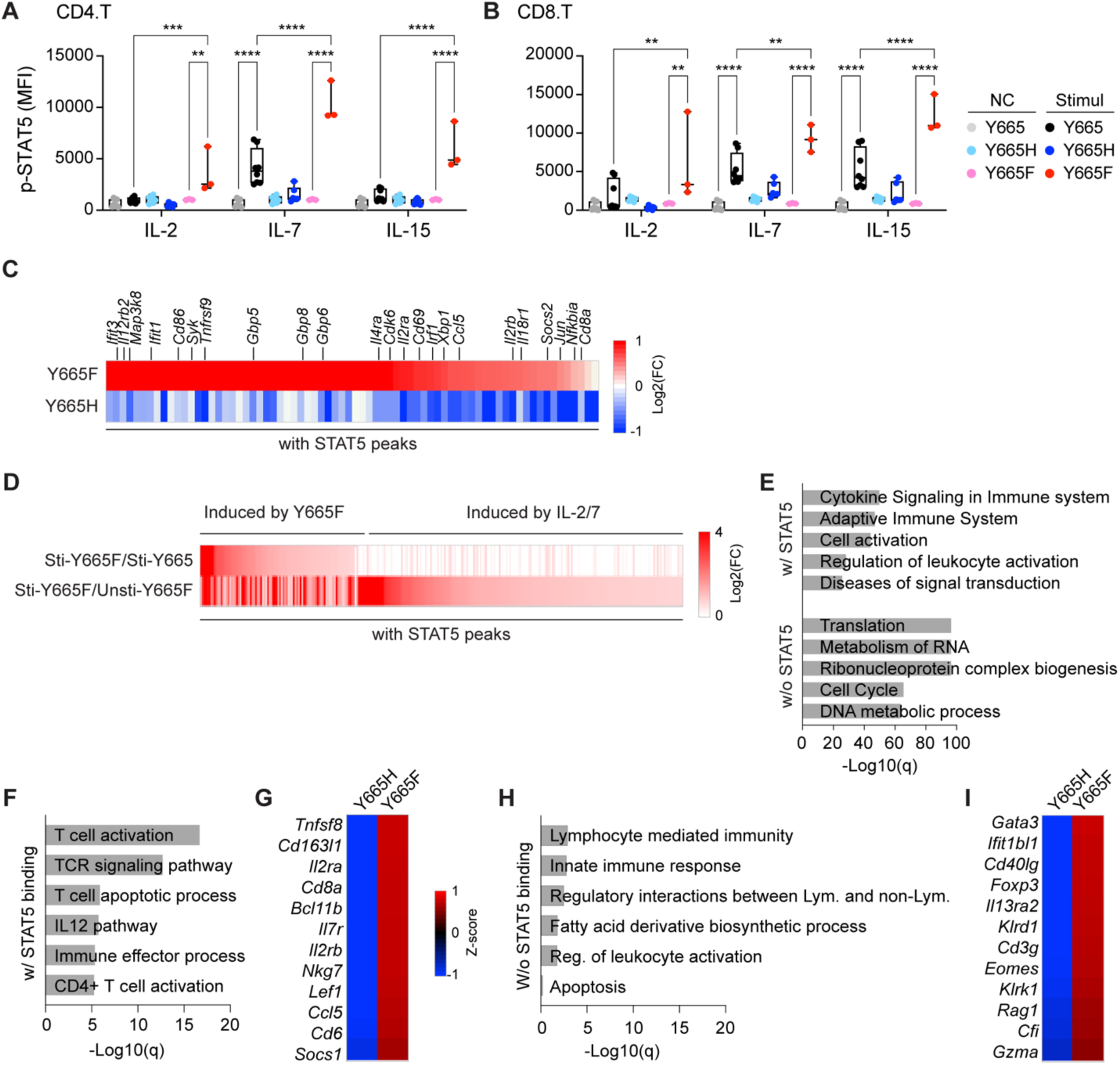
Opposing transcriptional activity of STAT5B^Y665F^ and STAT5B^Y665H^ at steady-state levels. **A-B** Quantification of phospho-STAT5 levels in CD4+ T cells (A) and CD8+ T cells (B) (Y665, *n* = 8; Y665H, *n* = 6; Y665F, *n* = 3). MFI, median fluorescent intensity. *P*-value are derived from two-way ANOVA with Tukey’s multiple comparisons test. \*\**P* < 0.01, \*\*\**P* < 0.0001, \*\*\*\**P* < 0.0001. **C** Heatmaps depicting fold changes of significantly regulated in *Stat5b^Y665F^* and *Stat5b^Y665H^*mice. **D** Heatmaps depicting fold changes of significantly upregulated and enriched genes in unstimulated or stimulated T cell from wild-type and *Stat5b^Y665F^* mice (IL-2/7 stimulated Y665, *n* = 4; IL-2/7 stimulated Y665F, *n* = 4; IL-2/7 stimulated Y665F, *n* = 3). **E** Gene categories expressed at significantly higher levels and depending on STAT5 binding in stimulated T cells from STAT5B^Y665F^ mice compared to WT mice. **F and H** Gene categories expressed at significantly higher levels and depending on STAT5 binding in STAT5B^Y665F^ mice compared to STAT5B^Y665H^ mice. **G and I** Heatmaps showing z-score of significantly upregulated and enriched genes in T cell activation (c) and apoptosis (e) between STAT5B^Y665H^ and STAT5B^Y665F^ mice (Y665H, *n* = 5; Y665F, *n* = 7).

### Opposing impact of STAT5B^Y665F^ and STAT5B^Y665H^ on transcriptional programs and enhancer establishment

To gain a more comprehensive understanding of the STAT5B^Y665F^ and STAT5B^Y665H^ mutants’ capacity to engage with the genome and activate biological programs we conducted transcriptomic and ChIP-seq studies. First, we performed ‘bulk’ RNA-seq on splenic T cells isolated from *Stat5b*^Y665F^ and wild-type mice following IL-2/IL-7 stimulation. Remarkably, 1010 DEGs with STAT5 binding on their regulatory elements were upregulated in *Stat5b*^Y665F^ mice relative to wild-type controls, with a particular enrichment in immune system processes and activation, but not in *Stat5b*^Y665H^ mice consistent with the level of p-STAT5 (Fig. 5C-E; Supplementary Table 2). While these *ex vivo* experiments demonstrated the enhanced capacity of the STAT5B^Y665F^ mutant to activate *bona fide* STAT5 target genes, this did not provide information on impact of the mutation on the steady state physiology of immune programs *in vivo*. To address the impact of the two mutants on steady state programs, we conducted bulk RNA-seq on T cells immediately upon their isolation, i.e. without additional interleukin stimulation. Expression of approximately 150 genes was elevated in *Stat5b*^Y665F^ splenocytes as compared to *Stat5b*^Y665H^ cells (Supplementary Table 3). The STAT5 bound genes were enriched in T cell activation and inflammatory response genes including genes typically induced by IFNs (Fig. 5F-I).

To identify those genes that are directly impacted by wild type and the two mutant STAT5B proteins, we conducted ChIP-seq on splenic T cells stimulated with IL-2 and IL-7, and measured STAT5B, PolII and H3K27ac occupancy. The data permitted the identification of candidate enhancers, both globally and at specific immune regulated loci (Fig. 6A-C). Approximately 4,400 STAT5B binding peaks coinciding with H3K27ac marks were identified in *Stat5b*^Y665F^ cells, while strikingly few sites bound by STAT5B were detected in *Stat5b*^Y665H^ cells, suggesting that this variant had little or no capacity to access the genome. In Stat5b^Y665F^ cells, STAT5B binding associated with H3K27ac marks at putative distal enhancers (1,465 genes), promoter regions (636 genes), intronic sequences (2,097 genes) and 3’UTR (229 genes) (Fig. 6B; Supplementary Table 4). Representative ChIP-seq data visualizing candidate enhancers located in the 5’ upstream region (*Cd8a*), promoter sequences (*Il18r1*), within introns (*Il2rb*) and 3’ flanking regions (*Il2rb*) are shown in Fig. 6D. We also identified 155 Stat5-bound enhancer clusters in Stat5b^Y665F^ cells (Fig. 6C; Supplementary Table 4), also referred to as super-enhancers or stretch enhancers. These enhancer clusters have been previously associated with pan-lineage STAT5 target genes, such as *Cish* and *Socs1*, and with lineage-specific genes, such as the *Il2ra (Li et al., 2017)* and the *Osm-Lif* locus. STAT5B binding was detected in the proximity of about 10% of the DEG in unstimulated splenic T cells and in approximately one half of the genes induced in primary splenic T cells cultured in the presence of interleukins (Supplementary Table 3), suggesting that they are direct transcriptional targets. This group includes genes encoding key immune receptors, including *Il2ra* and *Il12rb2*, and key transcription factors, including *Stat1*, *Irf1* and *Irf8* (Supplementary Table 3). These findings further emphasize that *Stat5b*^Y665F^ is a GOF mutation and STAT5B^Y665H^ is largely inactive.

**Fig. 6.**
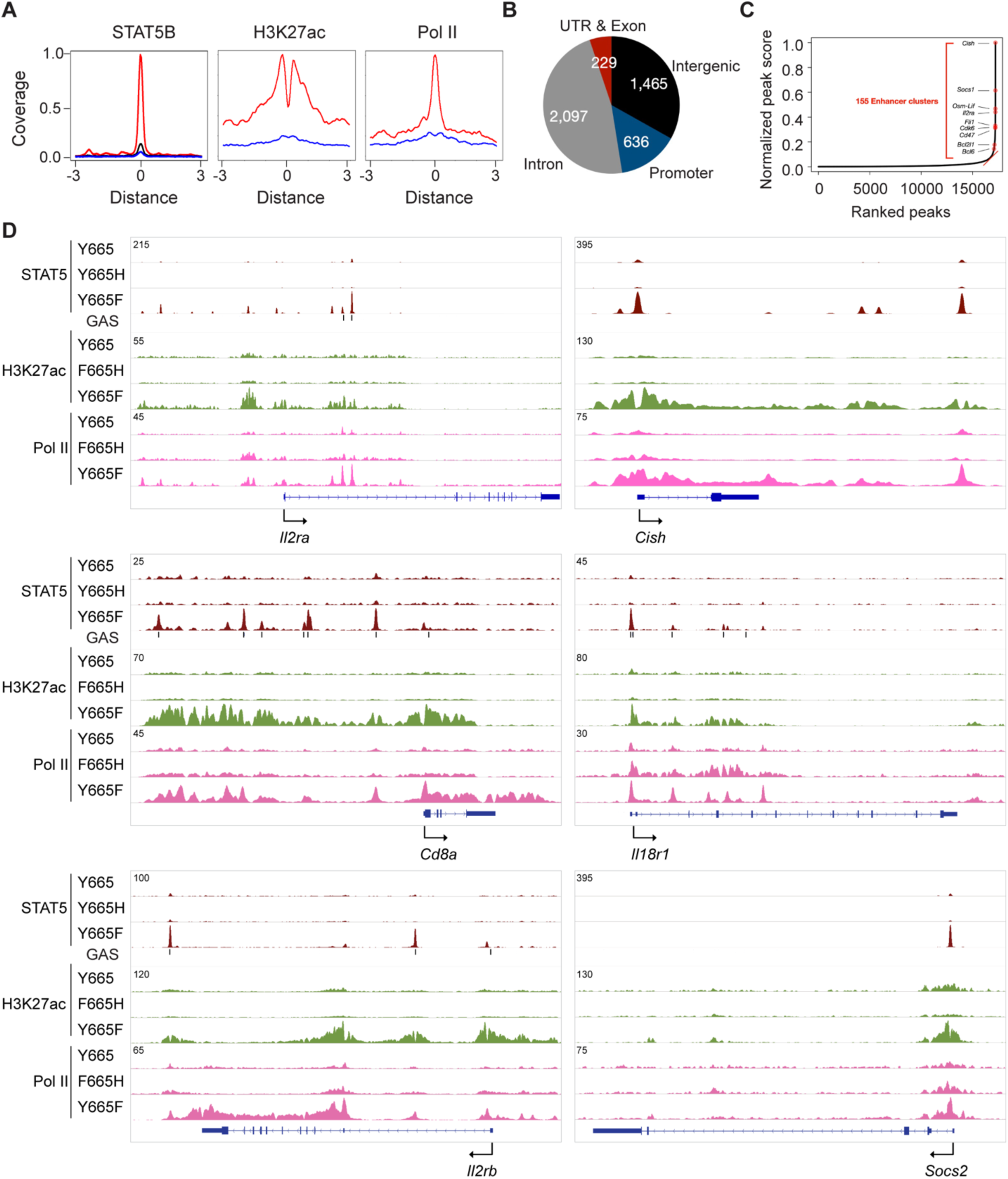
Inverse impact of STAT5B^Y665F^ and STAT5B^Y665H^ on the establishment of transcription enhancers and immune programs. **A** Coverage plots displaying the patterns of STAT5B binding, H3K27ac marks and Pol II loading on promoters of genes with or without STAT5B binding (blue, Y665H; red, Y665F; *n* = 2-3). **B** Distribution of STAT5B binding peaks across genomic regions. ChIP-seq signals within ± 3kbp of STAT5B binding regions identified in IL-2/IL-7 stimulated splenic T cells of Y665F mice are shown as pie chart. **C** Enhancer cluster analysis using IL-2/7 stimulated T cells of Y665F mice. Y-axis indicates the normalized peak score (enhancer cluster score) of STAT5B binding, whereas the x-axis represents the ranking of peaks. No enhancer clusters were identified in Y665H mice. **D** STAT5B binding, H3K27ac and Pol II loading at gene loci regulated by wild type and mutant STAT5B in IL-2/7 stimulated T cells.

## Discussion

The primary objective of this study was to understand the structural, genomic and pathophysiological impacts of two human leukemic STAT5B mutations (Kiel *et al*., 2014; Rajala *et al*., 2013), STAT5B^Y665F^ and STAT5B^Y665H^, identified in patients with T cell leukemias (Kiel *et al*., 2014; Rajala *et al*., 2013). For this we combined comprehensive *in silico* analyses, primary cell studies and experimental mouse genetics. The complementary *in silico* approaches, AlphaFold3, COORDinator, AlphaMissense and the pathogenicity prediction databases CADD (Rentzsch *et al*., 2019), REVEL (Ioannidis *et al*., 2016) and PolyPhen-2 (Adzhubei *et al*., 2013b) presented a complex picture of the mutations’ potential effects. While our structural analysis suggests both mutations could impact STAT5B dimerization, the inconsistencies of pathogenicity predictions highlight the challenges in relying solely on computational methods for assessing mutations in complex signaling pathways. *In vitro* transcriptomic studies using primary T cells confirmed gain-of-function (GOF) properties for STAT5B^Y665F^ and loss-of-function (LOF) for STAT5B^Y665H^, consistent with COORDinator predictions and PolyPhen-2 analysis. However, AlphaMissense, CADD and REVEL provided conflicting results. *Ex vivo* and *in vivo* studies in mice carrying these mutations further established STAT5B^Y665F^ as a GOF and STAT5B^Y665H^ LOF mutation. On a molecular level, STAT5B^Y665F^ exhibited an elevated capacity to establish genomic enhancers and hyperactivate transcriptional programs, leading to elevated CD8^+^ T cell populations. The elevation in Treg cells is compatible with previous observations of a permissive role for STAT5B in Treg development (Xiang *et al*, 2020). In contrast, STAT5B^Y665H^ failed to bind to genomic enhancers and activate interleukin-induced genetic programs, reminiscent to what has been observed in mice lacking Stat5b (Udy *et al*, 1997) and the STAT5B^Y699F^ mutant that fails to dimerize (Lin *et al*., 2024).

Even though both mutations studied have been identified in patients with T-PLL (Bhattacharya *et al*., 2022; Kiel *et al*., 2014; Kontro *et al*., 2014; Rajala *et al*., 2013) and have confirmed GOF activity *in vitro* (Pham *et al*., 2018), there was no evidence of overt leukemia in *Stat5b*^Y665H^ mice and *Stat5b*^Y665F^ mice, even at 11 months of age. However, some pathological features of STAT5B^Y665F^ mice, such as an increase of CD8^+^ and central memory cells, mirrors that observed in T-LGLL patients. Notably, the hyperactive STAT5B^Y665F^ galvanizes similar genetic programs in T-LGLL patients and in the mutant mice. Elevated expression of cytotoxicity-associated transcripts *Klrd1*, *Klrg1*, and *Gzma* is present in both patients (Huuhtanen *et al*, 2022; Rajala *et al*., 2013) and in mutant mice. We found that *Nkg7*, encoding the Natural Killer Cell Granule Protein 7, is a direct Stat5b target gene and highly induced in immune cells from *Stat5b*^Y665F^ mice. NKG7 is expressed in CD4+ and CD8+ T cells and has shown to be important in the inflammatory process (Lelliott *et al*, 2022; Ng *et al*, 2020). Although STAT5B^Y665F^ exerts transcriptional hyperactivity in humans and mice, it failed to directly induce T cell leukemia in mice, suggesting that the acquisition of additional secondary mutations is needed.

In light of human data (Kiel *et al*., 2014) and *in vitro* studies (Pham *et al*., 2018) it was surprising that STAT5B^Y665H^ was a LOF mutation and failed to activate genetic programs, both in our primary cell and mouse studies. AlphaFold3 predicted that the histidine substitution introduces a positively charged imidazole, potentially destabilizing the dimer is in full agreement with our findings. However, neither AlphaMissense, CADD, REVEL or PolyPhen2 predicted a LOF, highlighting the need for experimental validation to accurately determine the pathophysiological effects in an *in vivo* setting.

Congenital inactivating human STAT5B mutations, associated with severe growth failure and immune deficiencies, have been identified in several Laron syndrome patients(Hwa, 2021; Kofoed *et al*, 2003). Mutations such as STAT5B^A630P^ (Kofoed *et al*., 2003) and STAT5B^F646S^ (Scaglia *et al*, 2012) are autosomal recessive variants located within the SH2 domain, likely disrupting phosphate group binding on activated receptors or impairing STAT5 dimerization. Similar to STAT5B^Y665H^, these mutations fail to activate transcription and are associated with T-cell lymphopenia, a phenotype also observed in our mutant mouse models.

This study emphasizes the importance of adopting a multidisciplinary approach to investigate the effects of mutations associated with human diseases. While *in silico* tools like AlphaFold3 and COORDinator can predict structural impacts of mutations, functional algorithms such as AlphaMissense and PolyPhen-2, designed to assess pathogenicity, often produce inconsistent results. Similarly, *in vitro* cell culture studies provide only a limited perspective on the physiological and molecular consequences of mutations. A more comprehensive understanding of mutation pathogenicity can be achieved by integrating data from human patients with insights gained from experimental mouse genetics.

## Acknowledgements

We thank the NHLBI sequencing core for NGS. This work utilized the computational resources of the NIH HPC Biowulf cluster (http://hpc.nih.gov). H.K.L., S.L., P.A.F. and L.H. were supported by the Intramural Research Programs (IRPs) of National Institute of Diabetes and Digestive and Kidney Diseases (NIDDK), J.C, X.F., Z.W., C.L. and N.S.Y. were supported by Intramural Research Programs (IRPs) of National Heart, Lung, and Blood Institute (NHLBI), and R.P. and J.J.O. were supported by Intramural Research Programs (IRPs) of National Institute of Arthritis and Musculoskeletal and Skin Diseases (NIAMS). A.B.S. was supported by NIH Training grant T32AI162624. J.A.S. and F.B. were supported by the National Institute of General Medical Sciences of the NIH under R35 GM149227 to A.K.

## Contributions

Initial design of the study (LH, HKL, CL), further design input during the project (PAF, AVV, JC, XF, JJO, NSY, JAS, FB, AK), experimental processes (HKL, AVV, RLP, JC, FX, ZW, ABS, FB), data analysis and interpretation (HKL, SGL, JC, XF, LH, PAF, AVV, RP, MD, JAS, FB, AK), writing manuscript (HKL, LH, AVV and PAF).

## Methods

### Mice

All animals were housed and handled according to the Guide for the Care and Use of Laboratory Animals (8th edition) and all animal experiments were approved by the Animal Care and Use Committee (ACUC) of National Institute of Diabetes and Digestive and Kidney Diseases (NIDDK, MD) and performed under the NIDDK animal protocol K089-LGP-20. CRISPR/Cas9 and base editing targeted mice were generated using C57BL/6N mice (Charles River) by the Transgenic Core of the National Heart, Lung, and Blood Institute (NHLBI). Single-guide RNAs (sgRNA) were obtained Thermo Fisher Scientific’s In Vitro Transcription Service (Supplementary Table 5). Single-strand oligonucleotide donor was obtained from IDT (Supplementary Table 5). For the *Stat5b*^Y665H^ mutant mice, the ABE mRNA (50 ng/μl) and Y665H sgRNA (20 ng/μl) were co-microinjected into the cytoplasm of fertilized eggs collected from superovulated C57BL/6N female mice (Charles River Laboratories). For the *Stat5b*^Y665F^ mutant mice, a single-strand oligonucleotide donor contained the desired Y (TAC) to F (TTT) change and a silent C to G change. The silent mutation does not result in amino acid change but can destroy the sgRNA PAM and hence stopping Cas9 from further cutting after the oligo template was successfully knocked in. The *Stat5b*^Y665F^ sgRNA was first mixed with Cas9 protein (IDT) to form Cas9 RNP complex, which was co-electroporated with the oligo template into zygotes collected from C57BL/6N mice using a Nepa21 electroporator (Nepa Gene Co) following procedures described by Kenako (Kaneko, 2023). The microinjected or electroporated zygotes were cultured overnight in M16 medium (Millipore Sigma) at 37°C with 6% CO_2_. Those embryos that reached 2-cells stage of development were implanted into the oviducts pseudopregnant surrogate mothers (Swiss Webster mice from Charles River). All mice born to the foster mothers were genotyped by PCR amplification and Sanger sequencing (Quintara Biosciences) and automate genotyping using a TaqMan-based assay (Transnetyx) with genomic DNA from mouse tails.

Two-months old mice were used in the experiments. Tissues were collected from 2-month-old male mice and used immediately or stored at −80°C. Dermatitis was monitored until the age of 7-month-old and weights were measured until 10-week-old age.

### Whole exome sequencing and data analysis

Genomic DNA was isolated from tail tissue using Wizard Genomic DNA Purification Kit (Promega). Exome sequencing and bioinformatics analyses were performed at Psomage. Target capture for the exome was performed on each sample using SureSelect Mouse All Exon kit (Agilent Technologies). DNA was subjected to SureSelect Target Enrichment System for paired-end DNA library preparation. Whole exome sequencing was performed on a NovaSeq 6000 instrument (Illumina).

Sequencing reads were aligned to the mouse reference (mm10) using BWA 0.7.10. After excluding chimeric reads, the duplicated reads were eliminated using Picard. GATK3.v4 ‘IndelRealigner’ and ‘Table Recalibration’ were used for local realignment and for recalibrating the quality scores, respectively. For SNV/Indel calling in multi-sample analysis, GATK ‘HaplotypeCaller’ was used for comparison with the reference genome. For SNV calling in matched-pair analysis, ‘Selectvariants’ was used to compare the difference between WT and mutant mouse. Annotation for all variants was made using dbSNP142.

### Cell Counts and flow cytometry

Blood was collected from retro-orbital sinus into Eppendorf tubes in the presence of 5 mM ethylenediaminetetraacetic acid (EDTA, Sigma). Complete blood counts (CBC) were performed using an Element HT5 analyzer (Heska Corporation).

Mice were euthanized by CO_2_ to collect spleen, lymph node, tibia and femur. Spleens and lymph nodes were mechanically dissociated and filtered through a 70-micron strainer. Bone Marrow (BM) cells were extracted from bilateral tibiae and femurs, filtered through 85-micron nylon mesh and number counted by Vi-Cell counter (Beckman Coulter, Miami FL). Total BM cells per mouse were calculated based on the assumption that 2 tibiae and 2 femurs contain 25% of total BM cells. To stain cell surface antigens, peripheral blood, spleen and BM cells were first incubated in ACK buffer to remove red blood cells and were then stained with antibody mixtures for flow cytometry analysis.

After cell surface and fixable viability staining of single-cell suspensions with fluorophore-conjugated antibodies in PBS, intracellular staining was performed after fixing and permeabilizing cells with the Foxp3/Transcription Factor Staining kit (eBioscience) per manufacturer’s instructions. For measuring levels of phosphorylated STATs after *in vitro* stimulation, single-cell suspensions were stained with cell surface fluorophore-conjugated antibodies and fixable viability dye in PBS, fixed for 20 min at 4°C with BD Fix/Perm solution (Cat #554722), permeabilized with 100% cold methanol for 20min at 4°C, and stained with anti-p-STAT5 fluorophore-conjugated antibodies in 1xPerm/Wash Buffer (BD Biosciences, #557885) for 30 min at room temperature. The following antibodies were used for flow cytometry: BUV395-CD19 (ID3), BUV496-CD4 (GK1.5), BUV737-CD62L (MEL-14), BUV805-gdTCR (GL3), BV421-CD8a (53-6.7), efluor450-Foxp3 (FJK-16s), BV480-TCRb (H57-597), BV510-CD11b (M1/70), BV570-CD44 (IM7), BV605-CD25 (PC61), AF532-CD45 (30-F11), Spark Blue 550-Ly6G (1A8), PerCP-efluor710-NKp46 (29A1.4), AF647-pSTAT5 (C71E5), Zombie NIR, APC-NK1.1 (PK136), CD3, CD4, CD8, CD45R, Gr-1, CD11b, Ter119, CD117, Sca-1, CD48, CD150, CD34, CD16/32 (Biolegend). 7AAD and annexin V were purchased from BD Biosciences. Stained cells were acquired using BD FACSCanto II and BD LSR Fortessa flow cytometry operated by FACSDiva software (Becton Dickson) and 5-laser Cytek Aurora performed with FlowJo v10.

### T cell isolation and *In vitro* stimulation

For pSTAT5 level, lymphocytes isolated from spleen were stimulated at 37°C in complete RPMI alone or with 1000U/mL of recombinant mouse IL-2 (R&D systems), 10ng/mL of IL-7 (R&D systems), and/or 20ng of IL-15 (R&D systems) for 30 minutes.

For RNA-seq and ChIP-seq, total T cells from isolated splenocytes were isolated using EasySep Mouse T cell isolation kit (Stemcell, #19851) and cultured with recombinant mouse IL-2 and IL-7 from 24 hours.

### Western blot

Proteins (100 μg) from mouse mammary tissues were extracted with lysis buffer (50 mM Tris-Cl pH 8.0, 150 mM NaCl, 0.5% Na-DOC, 1% NP-40, 0.1% SDS, 5 mM EDTA, 1 mM PMSF, and protease inhibitor cocktail), separated on a 4-12% NuPage gradient gel (Invitrogen) and transferred to a PVDF membrane (Invitrogen). Membranes were blocked for 1 h with 5% nonfat dry milk in PBS-T buffer (PBS containing 0.1% Tween 20) and incubated for 1.5 hr at 4°C with the primary antibody against STAT5B (ThermoFisher scientific, 13-5300), phosphor-STAT5 (Abcam, ab32364) and GAPDH (Santa cruz, sc-47724). After washing, membranes were incubated for 1 hour with HRP-conjugated secondary antibodies (Cell signaling). Labeled protein bands were detected using an enhanced chemiluminescence system (Thermo scientific) and Amersham Imager 600 (GE healthcare).

### Histology

Isolated tissues from *mutant mice* were fixed with 10% neutral formalin solution and stored with 70% EtOH. Samples were processed for paraffin sections and stained with H&E at Histoserv.

### Isolation of Stat5a/b deficient CD4 T cells

The *Cd4-Cre* transgene was introduced into *Stat5a/b* floxed mice(Cui *et al*., 2004) resulting in the deletion of the *Stat5* locus in both CD4 and CD8 cells at the ‘double positive’ stage of thymic development. Starting population are sorted naive CD4+ T cells (live, CD4+CD44 lowCD25-neg) from poled lymph nodes and spleens. The sequenced population are sorted RV-transduced CD4+ T cells (live, CD4+GFP+). Cells were transduced with an empty retroviral vector or vectors encoding the native and mutant STAT5B isoforms, cultured for 48 hours with IL-2 and subsequently subjected to FACS analysis and RNA-seq.

### Total RNA sequencing (RNA-seq) and data analysis

Total RNA was extracted from frozen mammary and salivary tissue from wild-type and mutant mice and purified with RNeasy Plus Mini Kit (Qiagen, 74134). Ribosomal RNA was removed from 1 μg of total RNAs and cDNA was synthesized using SuperScript III (Invitrogen). Libraries for sequencing were prepared according to the manufacturer’s instructions with TruSeq Stranded Total RNA Library Prep Kit with Ribo-Zero Gold (Illumina, RS-122-2301) and paired-end sequencing was done with a NovaSeq 6000 instrument (Illumina).

Total RNA-seq read quality control was done using Trimmomatic (Bolger *et al*, 2014) (version 0.36) and STAR RNA-seq (Dobin *et al*, 2013) (version STAR 2.5.4a) using paired-end mode was used to align the reads (mm10). HTSeq(Anders *et al*, 2015) was to retrieve the raw counts and subsequently, R (https://www.R-project.org/), Bioconductor (Huber *et al*, 2015) and DESeq2 (Love *et al*, 2014) were used. Additionally, the RUVSeq (Risso *et al*, 2014) package was applied to remove confounding factors. The data were pre-filtered keeping only those genes, which have at least ten reads in total. Genes were categorized as significantly differentially expressed with log2 fold change >1 or <-1 and adjusted p-value (pAdj) <0.05 corrected for multiple testing using the Benjamini-Hochberg method were considered significant and then conducted gene enrichment analysis (Metascape, https://metascape.org/gp/index.html#/main/step1). The visualization was done using dplyr (https://CRAN.R-project.org/package=dplyr) and ggplot2 (Wickham, 2009).

### Single-cell RNA sequencing (scRNA-seq) and data analysis

Single-cell suspensions were then immediately loaded on the 10X Genomics Chromium Controller with a loading target of 10,000 cells. Libraries were generated using the Chromium Next GEM Single Cell 5′ Kit v2 (Dual Index) and Chromium Single Cell Mouse TCR Kit 5′ v2 (Dual Index) according to the manufacturer’s instructions. Libraries were sequenced using the NovaSeq 6000 instrument (Illumina).

The raw reads were aligned and quantified using the Cell Ranger with Feature Barcode addition (version 6.0, 10X Genomics) against the GRCm38 mouse reference genome. The quality control, normalization, dimension reduction, cell clusters, UMAP projection, and cell type annotation were performed using Seurat (version 4.3) (Hao *et al*, 2021).

### Chromatin immunoprecipitation sequencing (ChIP-seq) and data analysis

The frozen-stored tissues were ground into powder in liquid nitrogen. Chromatin was fixed with formaldehyde (1% final concentration) for 15 min at room temperature, and then quenched with glycine (0.125 M final concentration). Samples were processed as previously described (Metser *et al*, 2016). The following antibodies were used for ChIP-seq: STAT5A (Santa Cruz Biotechnology, sc-271542), STAT5B (R&D systems, AF1584; ThermoFisher scientific, 13-5300), H3K27ac (Abcam, ab4729) and RNA polymerase II (Abcam, ab5408). Libraries for next-generation sequencing were prepared and sequenced with the NovaSeq 6000 instrument (Illumina).

Quality filtering and alignment of the raw reads was done using Trimmomatic (Bolger *et al*., 2014) (version 0.36) and Bowtie (Langmead *et al*, 2009) (version 1.2.2), with the parameter ‘-m 1’ to keep only uniquely mapped reads, using the reference genome mm10. Picard tools (Broad Institute. Picard, http://broadinstitute.github.io/picard/. 2016) was used to remove duplicates and subsequently, Homer (Heinz *et al*, 2010) (version 4.9.1) and deepTools (Ramirez *et al*, 2016) (version 3.1.3) software was applied to generate bedGraph files and normalize coverage, separately. Integrative Genomics Viewer (Thorvaldsdottir *et al*, 2013) (version 2.3.98) was used for visualization. Coverage plots were generated using Homer (Heinz *et al*., 2010) software with the bedGraph from deepTools (Ramirez *et al*, 2014) as output. R and the packages dplyr (https://CRAN.R-project.org/package=dplyr) and ggplot2 (Love *et al*., 2014) were used for visualization. Each ChIP-seq experiment was conducted for two replicates and the correlation between the replicates was computed by Spearman correlation using deepTools.

To identify high-confident STAT5B binding peaks, we defined the STAT5B binding peaks which coincide with H3K27ac marks within ±500 bp and contain at least one GAS motif (5’-TTCNNNGAA-3’) within the peak region. The Integrative Genomics Viewer (IGV) tools were used for ChIP-seq signal visualization. Tag density plots and heatmaps were generated using computeMatrix and plotHeatmap tools implemented in deepTools (version 3.1.3). To examine enhancer clusters, we utilized findPeaks.pl with the following parameter settings: -style super-superSlope −1000. Enhancer clusters were identified as regions with “slope” (focus ratio)/(region size annotation enhancer) greater than 1. For further analyses, only protein-coding genes were used.

### Protein structure prediction and analysis

The STAT5B dimer structure was predicted using AlphaFold3 (https://alphafoldserver.com). The input sequences included human STAT5B (UniProt accession P51692) and the DNA fragment sequence (5’-GTTTC<u>TTC</u>TGA<u>GAA</u>GTACC-3’) derived from STAT5-bound Il2ra enhancer. The structure prediction was performed with pY699 phosphorylation parameter. Structural analysis and visualization were performed using PyMOL (version 2.4.1) and ChimeraX (version 1.8).

The dimeric structures of human STAT5A (Uniprot accession P42229, residues 620-715) and STAT5B (Uniprot accession P51692, residues 620-720) with phosphorylated residues STAT5A^pY694^ and STAT5B^pY699^ were modeled using AlphaFold3 (Abramson *et al*., 2024). The energetic consequences of amino-acid substitutions in STAT5A and STAT5B were predicted by COORDinator using these structures (Li *et al*., 2023). COORDinator mutation predictions were made by calculating the difference between the COORDinator-predicted energies (in arbitrary units) for the mutant and wild-type residues. COORDinator was fine-tuned to enforce a high correlation between predicted and experimental protein stability measurements for hundreds of thousands of mutations(Tsuboyama *et al*., 2023). The impact of a mutation on dimerization was predicted using the difference between the COORDinator energies for a residue in the absence vs. the presence of the dimerization partner, without modeling any change in conformation. Predicted effects of mutations on intra-molecular interactions with the C-terminal tail were obtained similarly, by treating the C-terminal tail as a separate chain. For example, the COORDinator-predicted energies for residues in the STAT5B C-terminal tail (residues 693-720) were computed in the context of the tail alone or in the context of the STAT5B monomer, using the conformations predicted for the STAT5B dimer. Similarly, energies for residues in the SH2 domain (residues 620-687) were computed in the presence vs. the absence of the C-terminal tail. When treating the STAT5B C-terminal tail as a separate chain, residues 688-692 were not included, to avoid effects from the amino- and carboxyl-groups of the artificially separated chains. The predicted importance of a residue was derived by summing the absolute values of the COORDinator energy predictions for all mutations at that site. For this analysis, we used code from COORDinator github (https://github.com/daveneff/Coordinator) and Terminator github (https://github.com/KeatingLab/terminator).

### *In silico* pathogenicity prediction

Multiple computational tools were employed to assess the potential pathogenicity of STAT5B variants. AlphaMissense (Cheng *et al*, 2023a) was used to predict the functional impact of mutations, with scores ranging from 0 (benign) to 1 (pathogenic). The Combined Annotation Dependent Depletion (CADD) (Rentzsch *et al*., 2019) tool was utilized to estimate the deleteriousness of variants, where PHRED scores above 20 indicate potentially damaging effects. The Rare Exome Variant Ensemble Learner (REVEL) (Ioannidis *et al*., 2016) scores, which range from 0 to 1, were calculated to assess the probability of pathogenicity. Additionally, PolyPhen-2 (Adzhubei *et al*, 2013a) analysis was performed to predict the possible impact of amino acid substitutions on protein structure and function, with scores ranging from 0 (benign) to 1 (probably damaging).

## Statistical analyses

All samples that were used for CBC, FACS and RNA-seq were randomly selected, and blinding was not applied. For comparison of samples, data were presented as standard deviation in each group and were evaluated with a 1-way or 2-way ANOVA multiple comparisons using PRISM GraphPad. Statistical significance was obtained by comparing the measures from wild-type or control group, and each mutant group. A value of \**P* < 0.05, \*\**P* < 0.001, \*\*\**P* < 0.0001, \*\*\*\**P* < 0.00001 was considered statistically significant. ns, no significant.

## Data availability

All data were obtained or uploaded to Gene Expression Omnibus (GEO). ChIP-seq, RNA-seq, and scRNA-seq data of wild-type and mutant tissues are under <u>GSE276308</u> (secure token: wlcremgmtvkfxuz), <u>GSE276311</u> (secure token: mlkxeassfvkbzcf) and <u>GSE276312</u> (secure token: clqlowasdvydpin). The fastq files and the processed *bedGraph* files can be downloaded from GEO (https://www.ncbi.nlm.nih.gov/gds/?term<u>=</u>) and imported into the IGV browser (https://software.broadinstitute.org/software/igv/download) with a reference genome (mm10).

## Supplementary Figures

**Supplementary Fig. 1.**
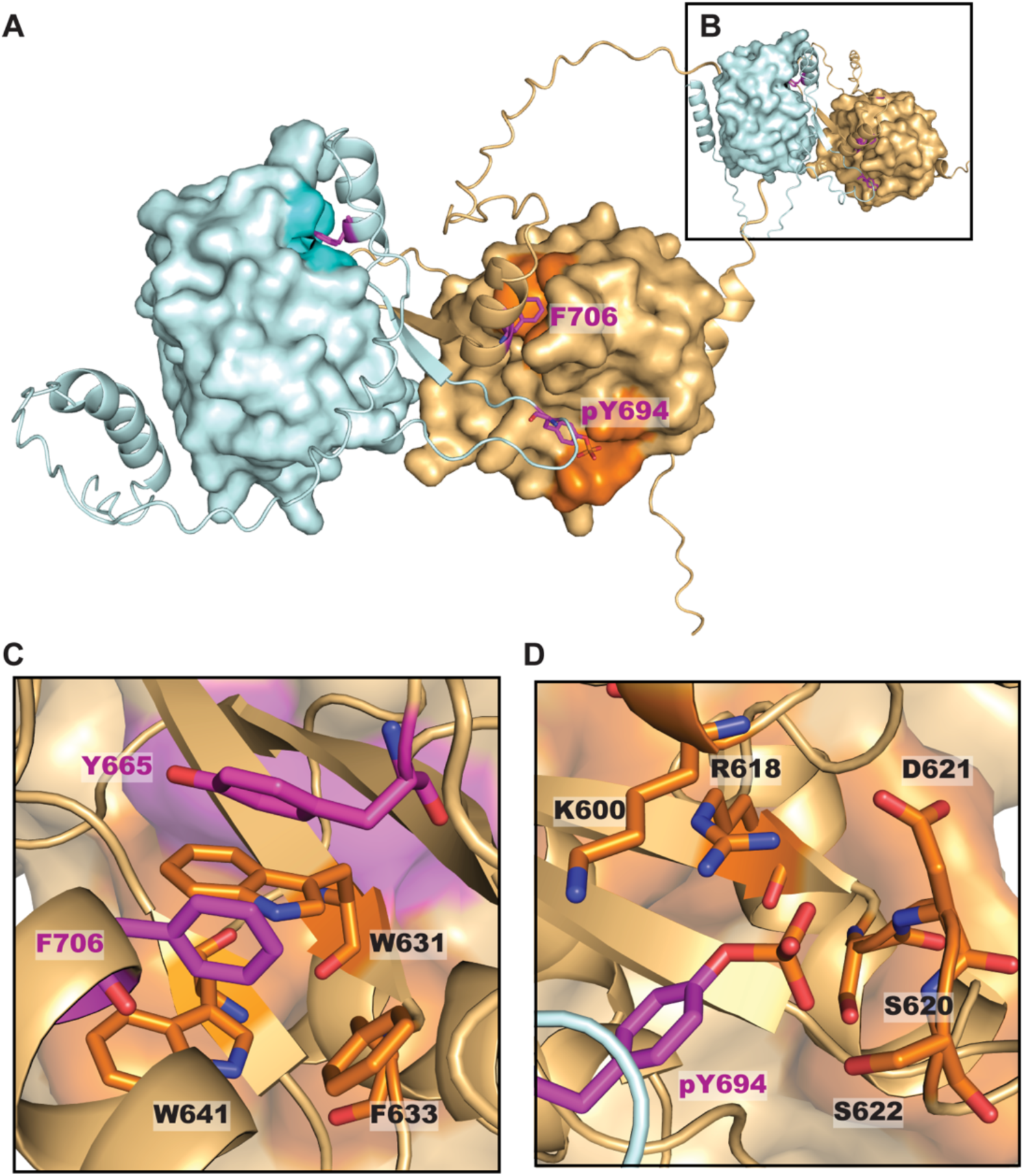
STAT5 SH2 dimerization modeled by AlphaFold3. **A** Structure of the human STAT5A SH2 homodimer generated by AlphaFold3. Binding pockets of key residues phospho-Tyr694 and Phe706 (purple), which are structurally analogous to phosphor-Tyr699 and Phe711 in STAT5B, are indicated. **B** Structure of the human STAT5B SH2 homodimer generated by AlphaFold3, which closely resembles the dimer interface modeled for the STAT5A homodimer. **C** The STAT5A model highlights the intramolecular interaction between Phe706 (purple) and the hydrophobic binding pocket (orange) containing key residue Tyr665 (purple). **D** The STAT5A model highlights the canonical SH2 docking interaction between the phospho-Tyr694 (purple) of the STAT5A C-terminal tail and the canonical SH2 binding pocket of a second STAT5A (orange).

**Supplementary Fig. 2.**
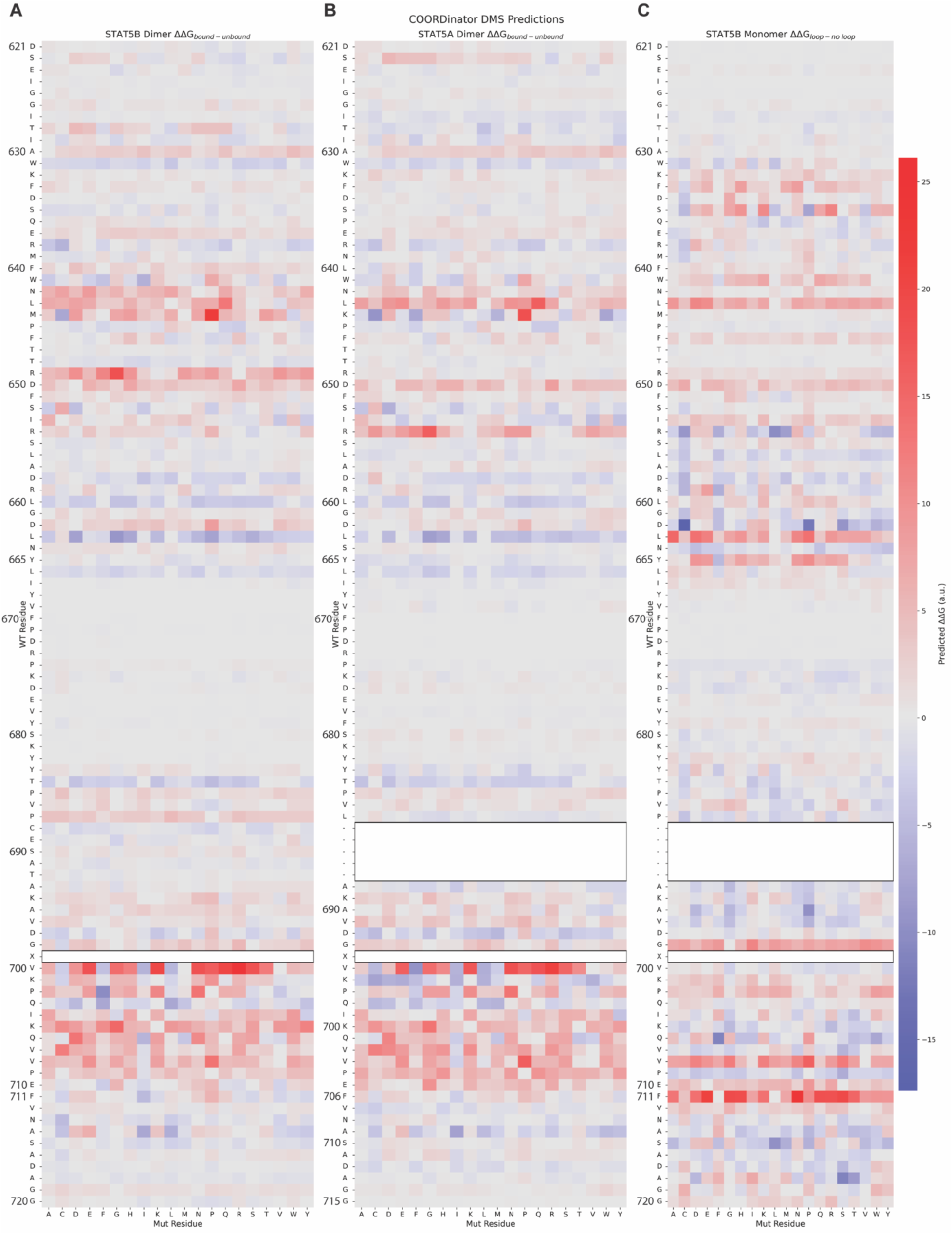
Energetic contributions of STAT5 SH2 domain residues to dimerization and intra-molecular interactions as predicted by COORDinator. Heat maps depicting the energetic consequences of amino-acid substitutions at each residue as predicted by COORDinator using AlphaFold3-generated structures. Relative mutational effects are annotated using arbitrary units with stabilizing mutations (-ΔΔG) depicted in blue and destabilizing mutations depicted in red (+ΔΔG). The intensity of the color corresponds to the extent of the change. Values for STAT5A^pY694^ and STAT5B^pY699^ are not included due to the inability of COORDinator to model post-translational modifications. **A** Energetic contributions of STAT5B residues to SH2 homodimerization. **B** Energetic contributions of STAT5A residues to SH2 homodimerization. Residues 688-692 are not present in STAT5A. **C** Energetic contributions of STAT5B residues to interaction with the C-terminal tail, when the tail is modeled as a separate chain. Residues 688-692 were not modeled to ensure the tail is properly seen by the model as a separate chain.

**Supplementary Fig. 3.**
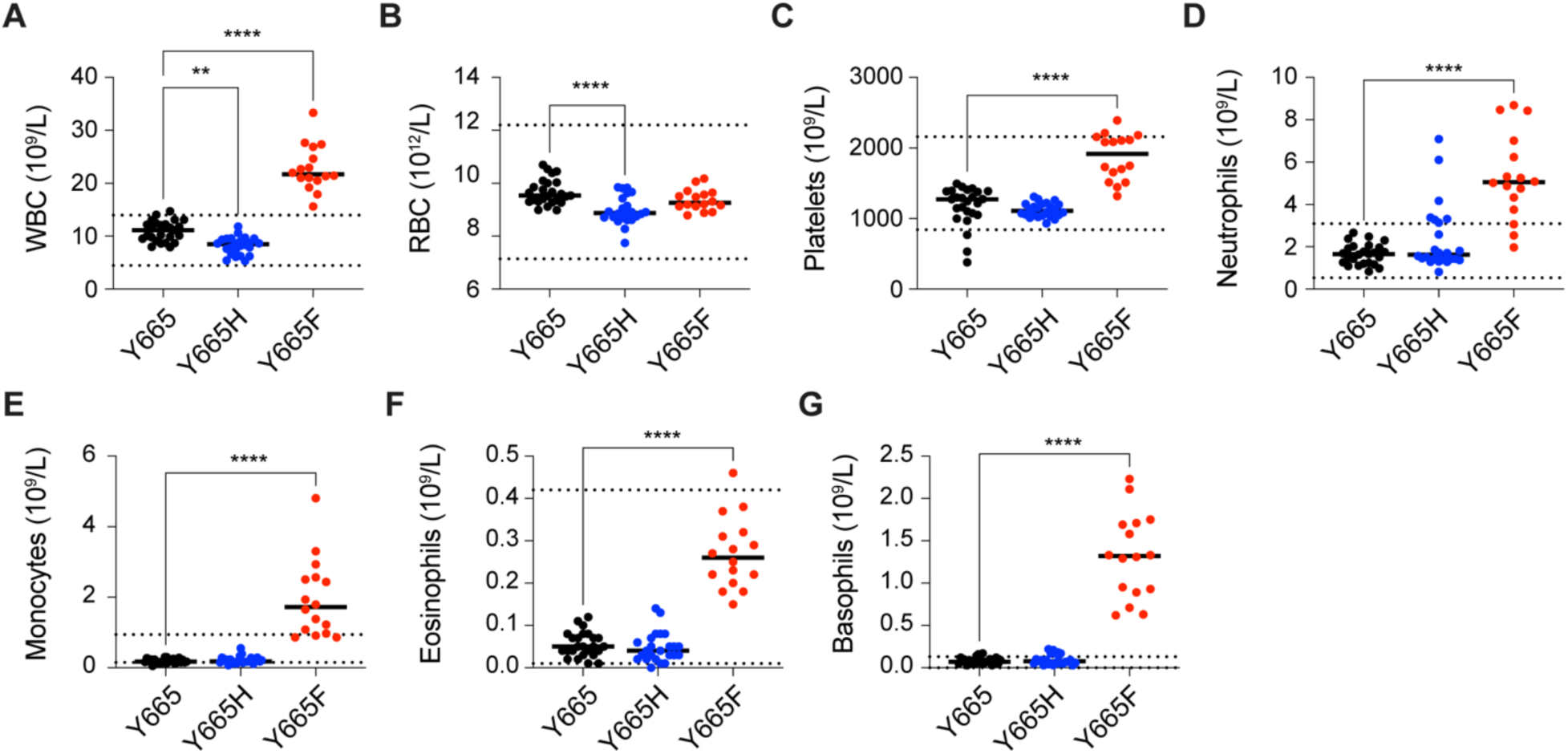
Hematological parameters and altered immune phenotypes of Stat5b mutant mice. Count of Blood Cells in peripheral blood from 7-10-week-old adult wild-type and mutant mice. Results are shown as the median of independent biological replicates (Y665, *n* = 25; Y665H, *n* = 24; Y665F, *n* = 16). *P*-value are from two-way ANOVA with Tukey’s multiple comparisons test. \**P* < 0.05, \*\**P* < 0.01, \*\*\**P* < 0.0001, \*\*\*\**P* < 0.0001.

**Supplementary Fig. 4.**
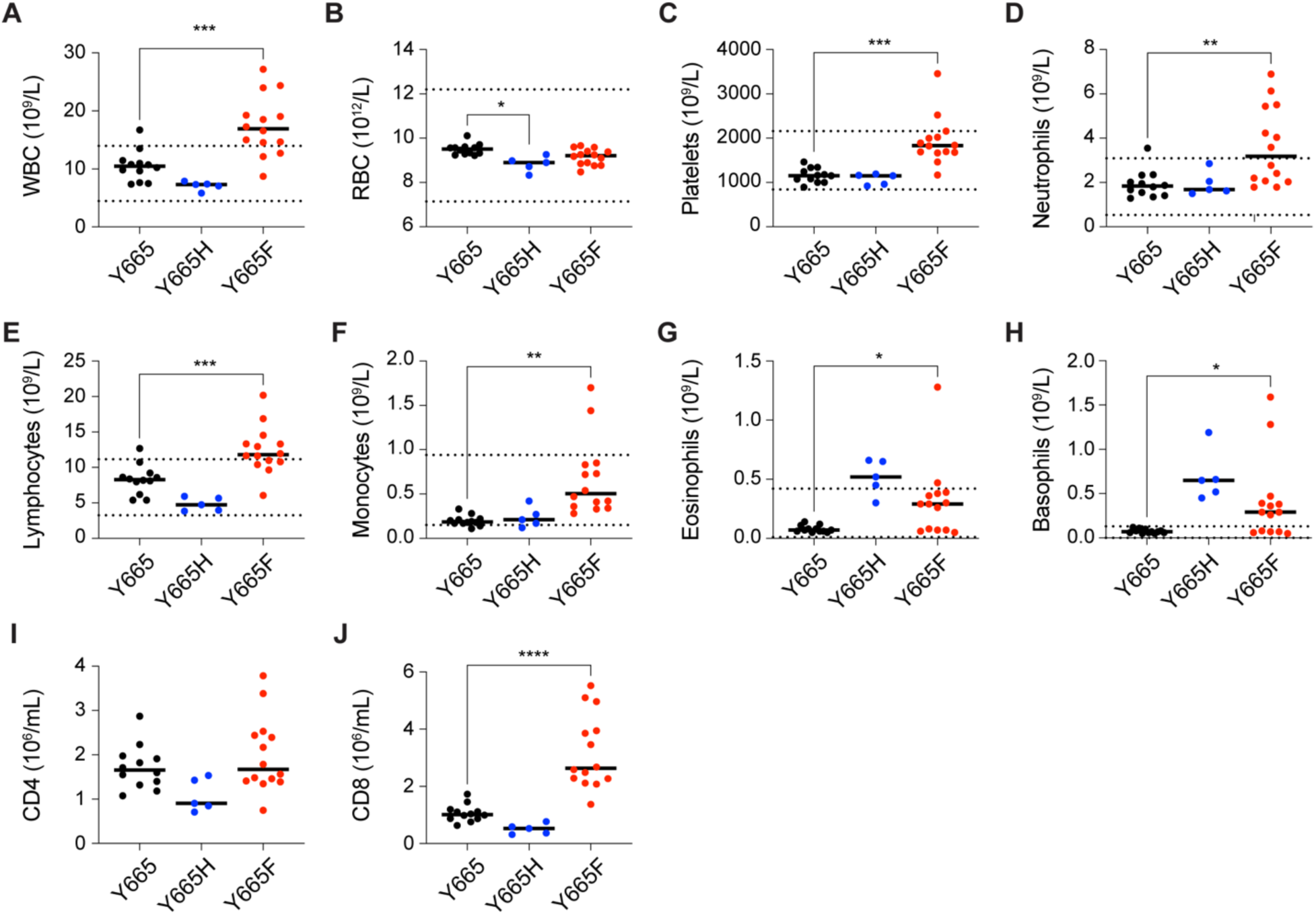
Hematological parameters and altered immune phenotypes of Stat5b mutant mice. A-H. Count of Blood Cells in peripheral blood from 11 months-old adult wild-type and mutant mice. Results are shown as the median of independent biological replicates (Y665, *n* = 12; Y665H, *n* = 5; Y665F, *n* = 14). **I-J** Numbers of subpopulation of immune cells identified by flow cytometry. Results are shown as the median of independent biological replicates (*n* = 5). Statistical significance was assessed using one-way ANOVA followed by Tukey’s multiple comparisons test. **n** Apoptosis rate detected by Annexin V and 7-amino-actinomycine staining using flow cytometry. *P*-value are from two-way ANOVA with Tukey’s multiple comparisons test. \**P* < 0.05, \*\**P* < 0.01, \*\*\**P* < 0.0001, \*\*\*\**P* < 0.0001.

## Supplementary Tables

**Supplementary Table 1.** List of genes that are significantly up-regulated by Stat5b mutant plasmids in CD4+ T cells from STAT5-deficient mice.

**Supplementary Table 2.** List of significantly up-regulated genes in IL-2/7 stimulated T cells from spleen of STAT5B^Y665F^ and STAT5B^Y665H^ compared to IL-2/7 stimulated T cells from Y665 and unstimulated T cells from STAT5B^Y665F^ with normalized read counts at spleen tissue, log2 (fold change), their *p*-value and adjusted *p*-value. List of genes with or without STAT5 binding on their regulatory elements.

**Supplementary Table 3.** List of significantly up-regulated genes in STAT5B^Y665F^ compared to STAT5B^Y665H^ with normalized read counts at spleen tissue, log2 (fold change), their *p*-value and adjusted *p*-value. List of genes with or without STAT5 binding on their regulatory elements.

**Supplementary Table 4.** List of STAT5B binding peaks, STAT5B bound enhancers and enhancer clusters in STAT5B^Y665H^ and STAT5B^Y665F^ mice.

**Supplementary Table 5.**
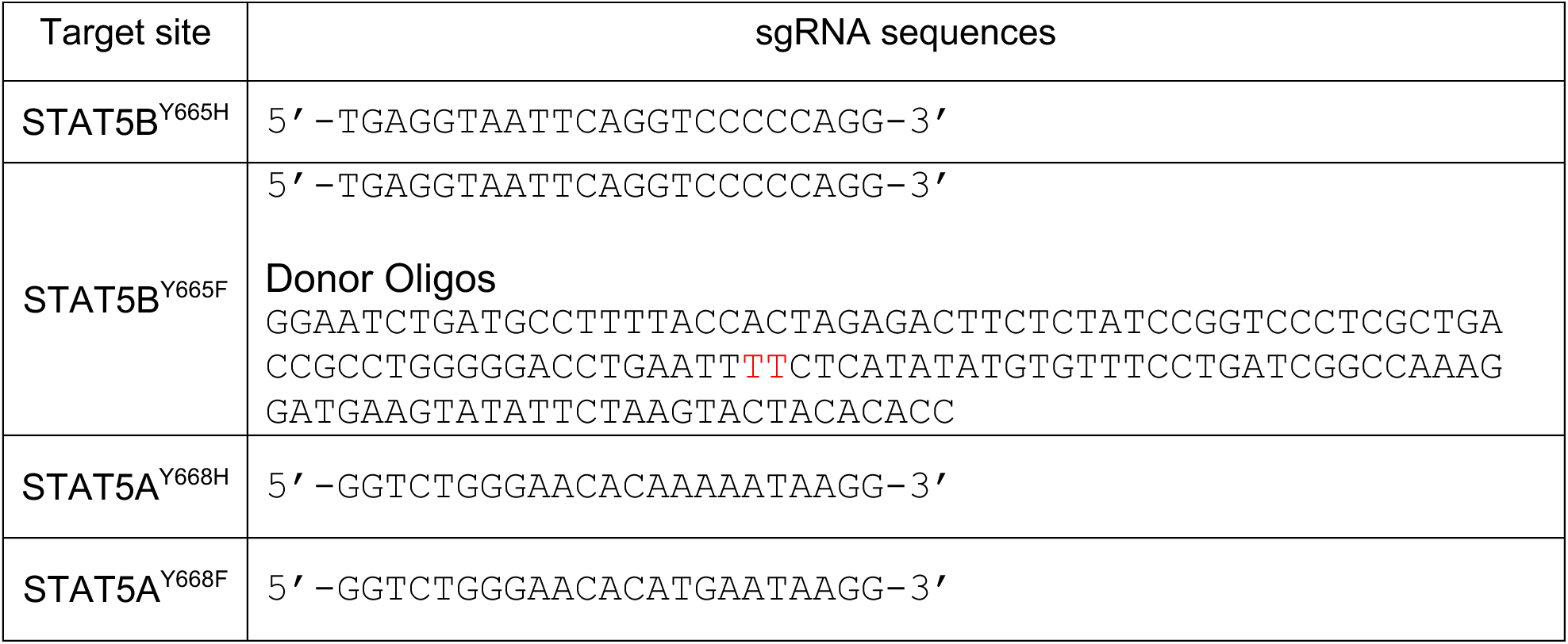
Sequences of sgRNA for CRISPR/Cas9 and base-editing targeted mice. The donor oligo is contained the desired Y (TAC) to F (TTT) change.

